# Real-time structural dynamics of late steps in bacterial translation initiation visualized using time-resolved cryogenic electron microscopy

**DOI:** 10.1101/390674

**Authors:** Sandip Kaledhonkar, Ziao Fu, Kelvin Caban, Wen Li, Bo Chen, Ming Sun, Ruben L. Gonzalez, Joachim Frank

## Abstract

Bacterial translation initiation entails the tightly regulated joining of the 50S ribosomal subunit to an initiator transfer RNA (fMet-tRNA^fMet^)-containing 30S ribosomal initiation complex (IC) to form a 70S IC that subsequently matures into a 70S elongation-competent complex (70S EC). Rapid and accurate 70S IC formation is promoted by 30S IC-bound initiation factor (IF) 1 and the guanosine triphosphatase (GTPase) IF2, both of which must ultimately dissociate from the 70S IC before the resulting 70S EC can begin translation elongation^1^. Although comparison of 30S^2–6^ and 70S^5,7–9^ IC structures have revealed that the ribosome, IFs, and fMet-tRNA^fMet^ can acquire different conformations in these complexes, the timing of conformational changes during 70S IC formation, structures of any intermediates formed during these rearrangements, and contributions that these dynamics might make to the mechanism and regulation of initiation remain unknown. Moreover, lack of an authentic 70S EC structure has precluded an understanding of ribosome, IF, and fMet-tRNA^fMet^ rearrangements that occur upon maturation of a 70S IC into a 70S EC. Using time-resolved cryogenic electron microscopy (TR cryo-EM)^10^ we report the first, near-atomic-resolution view of how a time-ordered series of conformational changes drive and regulate subunit joining, IF dissociation, and fMet-tRNA^fMet^ positioning during 70S EC formation. We have found that, within ~20–80 ms, rearrangements of the 30S subunit and IF2, uniquely captured in its GDP•P*_i_*-bound state, stabilize fMet-tRNA^fMet^ in its intermediate, ‘70S P/I’, configuration^7^ and trigger dissociation of IF1 from the 70S IC. Within the next several hundreds of ms, dissociation of IF2 from the 70S IC is coupled to further remodeling of the ribosome that positions fMet-tRNA^fMet^ into its final, ‘P/P’, configuration within the 70S EC. Our results demonstrate the power of TR cryo-EM to determine how a time-ordered series of conformational changes contribute to the mechanism and regulation of one of the most fundamental processes in biology.

---

Translation initiation is a fundamental step in gene expression that is essential for the overall fitness and viability of cells. In bacteria, the dynamic initiation reaction is kinetically controlled by three IFs (IF1, IF2, and IF3), which collaborate to ensure accurate selection of fMet-tRNA^fMet^ and its pairing with the mRNA start codon^11–14^. Canonical initiation begins with assembly of the 30S IC, followed by IF2-catalyzed joining of the 50S subunit to the 30S IC to form a 70S IC and maturation of the 70S IC into a 70S EC^1,15,16^. Given the essential nature of this process, structural intermediates formed during initiation in bacteria represent promising targets for the development of next-generation antibiotics^4,17,18^.

Ensemble rapid kinetic and single-molecule studies have led to the identification and characterization of several intermediate steps during the late stages of initiation. These studies have shown that subunit joining triggers rapid GTP hydrolysis by IF2^1,15,19–21^, dissociation of the IFs^1,13,19^, transition of the ribosomal subunits into their non-rotated inter-subunit orientation^22,23^, and accommodation of fMet-tRNA^fMet^ into the peptidyl-tRNA binding (P) site of the peptidyl transferase center (PTC)^24^. In addition, structures of various 30S^2–6^ and 70S ICs^5,7–9^ obtained by cryogenic electron microscopy (cryo-EM) have revealed intermediate ICs that vary in the conformation of the ribosome, IFs, and fMet-tRNA^fMet^. Nonetheless, notable discrepancies in the inter-subunit orientation of the ribosome and position of fMet-tRNA^fMet^ in several of the available 70S IC structures have made it difficult to arrive at a consensus structural model for initiation^5, 7–9^. Furthermore, the 70S ICs represented by the available structures were formed using a 70S ribosome and an IF2 bound to a non-hydrolyzable GTP analog (*e.g.*, GDPNP) that results in a biochemically trapped 70S IC, rather than by mixing the 50S subunit with a 30S IC that carries a native, GTP-bound IF2 and results in the formation of a 70S IC, which subsequently matures into a 70S EC. Consequently, the available 70S IC structures do not provide information about how the various structural intermediates that have been observed evolve over the course of the initiation reaction. Therefore, these structural studies have been unable to distinguish on-pathway intermediates formed during canonical initiation from spurious, off-pathway intermediates.

To circumvent this problem, and to capture authentic, on-pathway intermediates that are created during canonical translation initiation, we have employed mixing-spraying-based TR cryo-EM^10,25–28^. Previously, we have used this TR cryo-EM method to study the association of vacant 30S- and 50S subunits to form 70S ribosomes^25^, as well as to visualize transient structural intermediates formed during the ribosome recycling process^26^. In the current study, we have used mixing-spraying-based TR cryo-EM to investigate the IF-catalyzed joining of the 30S IC with the 50S subunit to form an authentic 70S IC that matures into a 70S EC. Using this approach, we have visualized, in real time and with near-atomic spatial resolution, the conformational rearrangements of the 30S and 70S ICs that promote and control subunit joining, IF dissociation, and fMet-tRNA^fMet^ positioning during 70S EC formation.

30S ICs were assembled by combining 30S subunits, mRNA, fMet-tRNA^fMet^, IF1, and IF2 with GTP in our optimized Tris-Polymix Buffer^29^. Previous ensemble rapid kinetic- and singlemolecule studies have shown that IF3 dissociates prior to^11^, or concomitant^13,30^ with, subunit joining in order to allow stable formation of the 70S IC and its maturation into the 70S EC^16,19,30^. Thus, in order to ensure that formation of the 70S IC would go to completion, IF3 was not included in the 30S IC. We began by determining whether the 30S IC was stable enough to maintain its integrity during injection into the mixing chamber and reaction channel of the mixing-spraying microfluidic chip as well as during spraying onto the EM grid. For this control experiment, the 30S IC was injected and mixed with an equal volume of Tris-Polymix Buffer lacking 50S subunits, using the microfluidic chip designed to give the longest reaction time (~600 ms), and subsequently sprayed onto an EM grid that was rapidly plunged into liquid ethane. The results of this control experiment confirmed that ~75 *%* of the 30S ICs remain intact during the mixing-spraying process (Extended Figure 1).

Ensemble rapid kinetic studies suggest that transient intermediates formed during initiation are populated on the sub-second timescale^11,13,15,19–21,31^. Using published rate constants^19,31^, we developed a kinetic model and analyzed how the populations of the expected structural species were predicted to vary as a function of time during subunit joining reactions in which 0.6 μM 50S subunits were mixed with 1.2 μM 30S ICs (Methods and Extended Figure 2). The analysis predicts that the population of 70S ICs carrying a GTP- or GDP•P*_i_*-bound IF2 is maximized at ~150 ms and that joining of the 50S subunit to the 30S IC to form a mature 70S EC is ~65 % complete within 600 ms. Using a set of microfluidic chips designed^27^ to provide reaction times of ~20 ms, ~80 ms, ~200 ms, and ~600 ms in our mixing-spraying TR cryo-EM apparatus (Extended Figure 3), we therefore mixed 0.6 μM of 50S subunits with 1.2 μM of 30S ICs and collected images at each time point. At each time point, two-dimensional (2D) classification of the images yielded 30S subunit-like, 50S subunit-like, and 70S ribosome-like particle classes. Subsequently, the particles from the ~20 ms, ~80 ms, ~200 ms, and ~600 ms time points were combined into a single dataset of 30S subunit-like, 50S subunit-like, and 70S ribosome-like particle and subjected to 3D classification (Methods and Extended Figure 4). 3D classification yielded the structures of five distinct classes: (1) a complex containing the 30S subunit, mRNA and fMet-tRNA^fMet^, but lacking IF1 and IF2; (2) the 30S IC; (3) the 50S subunit; (4) the 70S IC; and (5) the 70S EC. Notably, the populations of the 50S subunit, 70S IC, and 70S EC qualitatively followed the predicted kinetics (Figure1a and Extended Figure 2), with the population of the 50S subunit decreasing as the population of the 70S EC increases from ~20 ms to ~600 ms (Extended Figure 2). Among the five particle classes that we obtained, we selected the 30S IC, 70S IC, and 70S EC for further structural analysis (Figures 1b, 1c, and Figure 2a). To our knowledge, the 70S IC structure represents the first reported structure of a 70S IC obtained by mixing the 50S subunit with a 30S IC carrying a native, GTP-bound IF2, and the 70S EC structure represents the first reported structure of an authentic 70S EC. The resolutions of the 30S IC, 70S IC, and 70S EC were estimated to be 4.2 Å, 4.0 Å, and 3.9 Å, respectively, according to a resolution-estimating protocol that avoids overfitting and uses the Fourier shell correlation (FSC) with the 0.143 criterion^32^ (Extended Figure 5). Molecular Dynamics Flexible Fitting (MDFF)^33^ was then used to generate structural models of the 30S IC and 70S IC, while rigid-body fitting of previously published structures of the 30S and 50S subunits (PDB IDs: 2AVY and 2AW4, respectively) was used to generate a structural model of the 70S EC (see Methods).

**Figure 1:**
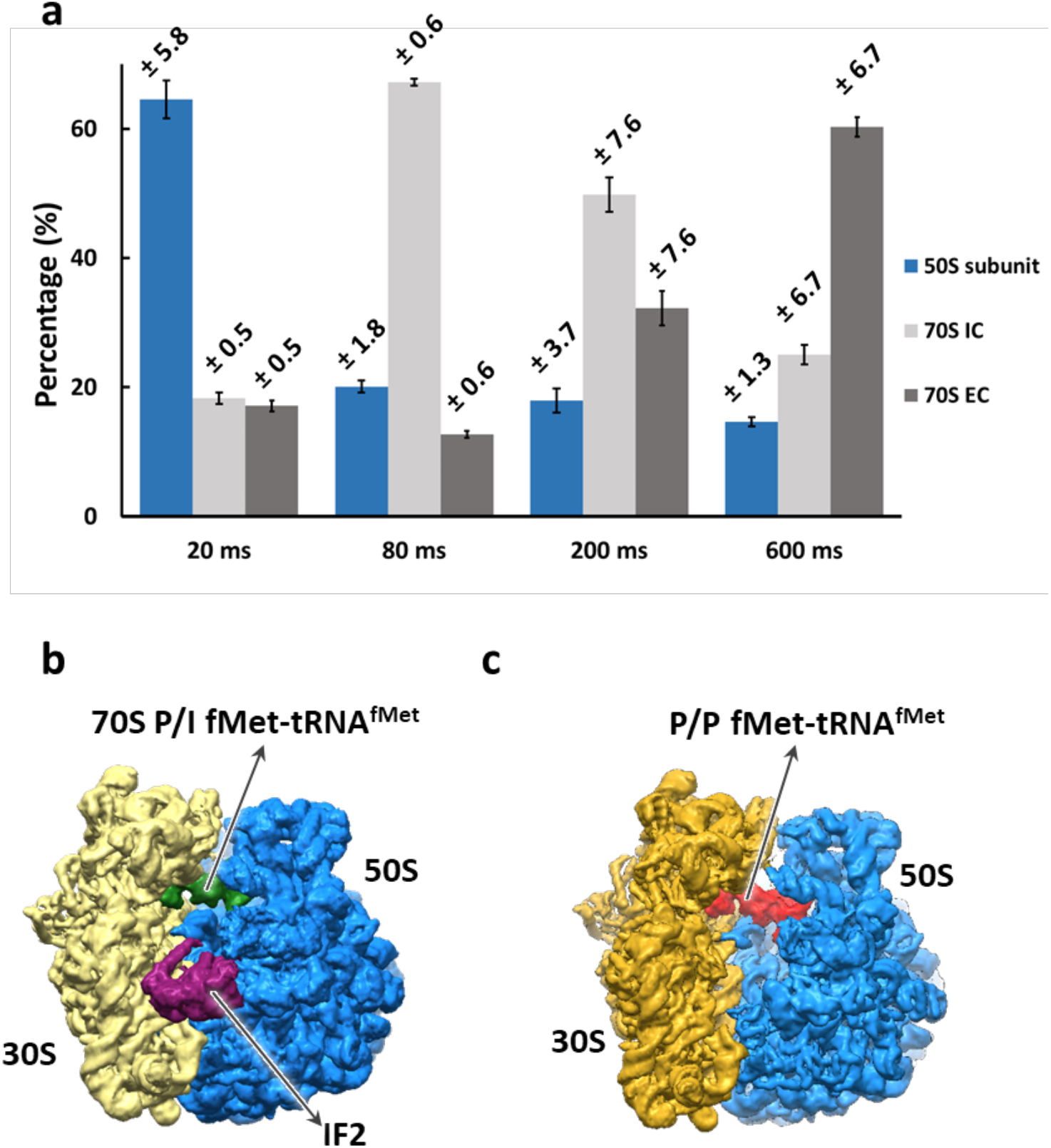
Structural and time-resolved population analyses of the 50S subunit, 70S IC, and 70S EC. **(a)** The populations of the 50S subunit, 70S IC, and 70S EC at the 20 ms, 80 ms, 200 ms, and 600 ms time points as obtained after 3D classification of the imaged particles. The error bars represent standard deviations obtained by repeating the 3D classification procedure three times for each time point. **(b-c)** The cryo-EM-derived Coulomb potential maps^34^ of the (b) 70S IC and (c) 70S EC.

**Figure 2:**
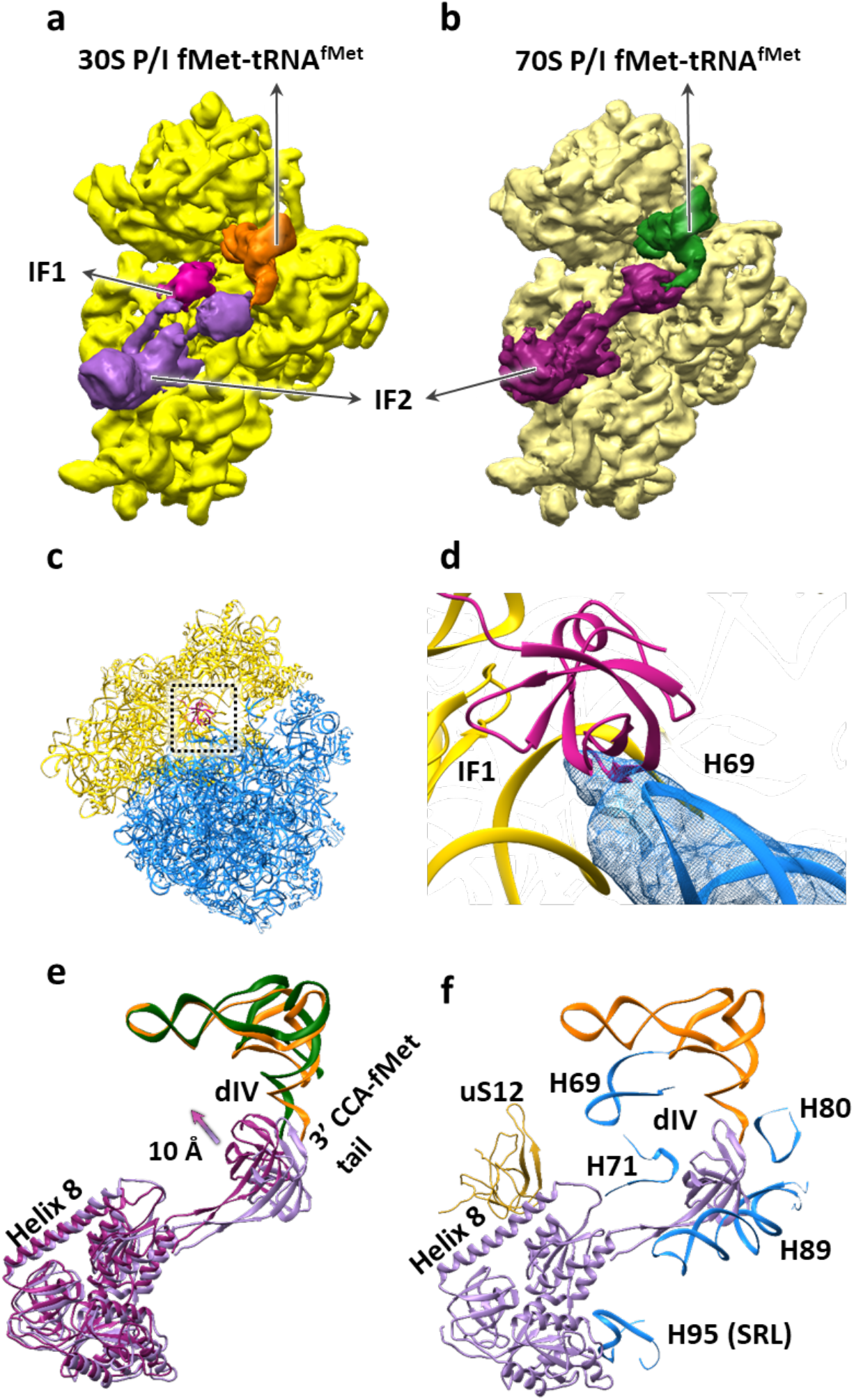
Ribosome, IF, and fMet-tRNA^fMet^ dynamics during 70S IC formation. **(a-b)** Cryo-EM-derived Coulomb potential maps^34^ viewed from the inter-subunit faces of (a) the 30S IC (with the 30S subunit shown in yellow) and (b) the 30S subunit (pale yellow), IF2, and fMet-tRNA^fMet^ from the 70S IC. **(c)** Superposition of the 30S subunits of the 30S IC and the 70S IC and analysis of the conformations of the 30S subunit and IF1 (shown in magenta) from the 30S IC and the 50S subunit from the 70S IC. The analysis reveals that rapid dissociation of IF1 upon 50S subunit joining to the 30S IC relieves a potential steric clash between IF1 and the 50S subunit that would take place during 70S IC formation. **(d)** A magnified view of the superposition shown in panel (c) highlights the potential steric clash between turn 1 of IF1 and H69 of the 50S subunit. **(e)** Superposition of the 30S subunits from the 30S IC and the 70S IC and comparative analysis of the conformations of IF2 and fMet-tRNA^fMet^ from the 30S IC (light purple and orange, respectively) and IF2 and fMet-tRNA^fMet^ from the 70S IC (dark purple and green, respectively). The analysis reveals that dIV of IF2 moves towards the inter-subunit face of the 30S subunit by ~10 Å and, as it rearranges from its 30S P/I to its 70S P/I configuration, the central domain and 3’ CCA-fMet tail of fMet-tRNA^fMet^ move slightly towards tRNA exit (E) site of the 30S subunit upon 50S subunit joining to the 30S IC and formation of the 70S IC. **(f)** Superposition of the 30S subunits from the 30S IC and the 70S IC and analysis of the conformations of IF2 from the 30S IC and uS12 (shown in pale yellow) of the 30S subunit of the 70S IC and H69, H71, H80, H89, and H95 (the sarcin-ricin loop (SRL)) (shown in blue) of the 50S subunit of the 70S IC that interact with IF2. The analysis reveals that the IF2 rearrangements shown in panel (e) relieve a potential steric clash between dIV of IF2 and H89 that would take place during 70S IC formation.

Analysis of the 70S ICs that are formed within the first ~20-80 ms after mixing 50S subunits with 30S ICs shows that all inter-subunit bridges are formed. Moreover, we find that IF1 has also dissociated from these 70S ICs (compare Figures 2a and 2b). This observation is significant because IF1 occupies a binding site between the cleft of 16S ribosomal RNA (rRNA) helix (h) 44, h18, and ribosomal protein uS12 on the 30S subunit that enables turn 1 of IF1, comprised of residues 18-21, to establish contacts with the minor groove of h44. Consequently, dissociation of IF1 relieves a strong steric clash that would otherwise exist between turn 1 of IF1 and 23S rRNA helix (H) 69 of the 50S subunit (Figure 2c and 2d). Because inter-subunit bridge B2a is formed by an interaction between h44 and H69, dissociation of IF1 early during subunit joining enables this critically important inter-subunit bridge to be established rapidly during initial 70S IC formation.

By the time ~80 ms has elapsed, the population of the 70S IC has reached its maximum and by ~200 ms, IF2 has dissociated from a significant fraction of this 70S IC population, resulting in the formation of mature 70S ECs, a process that continues through the 600 msec time point and beyond. Interestingly, the 70S IC that is captured in this study by mixing 50S subunits with 30S ICs carrying native, GTP-bound IF2 is in a semi-rotated inter-subunit orientation that is very similar to the orientation observed in the 70S IC reported by Allen and coworkers^7^ and the orientation of the major population of the 70S IC reported by Sprink and coworkers (*i.e.*, 70S-IC II)^9^. 70S IC-bound IF2 establishes three sets of interactions with the ribosome that help lock the 70S IC in the semi-rotated inter-subunit orientation. Specifically, helix 8 of IF2 interacts with the inter-subunit surface of uS12 in the 30S subunit; domain IV (dIV) of IF2 interacts with H69, H71, H89, and the loop-containing residues 77-85 of ribosomal protein uL16 of the 50S subunit; and dI of IF2 interacts with the sarcin-ricin loop (H95) of the 50S subunit (Extended Figure 6). Because the semi-rotated inter-subunit orientation of the 70S ribosome uniquely facilitates the simultaneous formation of these three sets of IF2-ribosome interactions, IF2 selectively stabilizes the 70S IC in this orientation.

Comparative analysis of the 30S and 70S ICs reveals subunit joining-dependent conformational changes of IF2 that facilitate formation of the 70S IC. Relative to its position on the 30S IC, dIV of IF2 is stabilized in a position that is ~10 Å closer to the 30S subunit, a structural transition that eliminates a potential steric clash with H89 (Figure 2e and 2f). Furthermore, the relatively early dissociation of IF1 during 70S IC formation increases the conformational freedom of helix 8 of IF2, allowing it to acquire a position that is closer to the 50S subunit (Figure 2e). Based on previous ensemble rapid kinetic studies^15,19–21^ demonstrating that the rate of GTP hydrolysis is relatively fast and nearly indistinguishable from the rate of initial subunit association, and that the rate of P*_i_* release is slower than the rate of IF2 dissociation, we propose that we have uniquely captured the native, GDP•P*_i_*-form of IF2 on the 70S IC. This proposal is supported by a structural analysis demonstrating that GDP•P*_i_* more precisely models the Coulomb potential map^34^ of the guanosine nucleotide bound to IF2 in the 70S IC than GTP does (Extended Figure 7). The similarity between the conformation of the GDP•P*_i_*-form of IF2 captured here and the non-hydrolyzable GTP analog-form of IF2 reported in all of the other 70S IC structures that have been published^5,7–9^ suggests that when IF2 hydrolyzes GTP, it does not immediately undergo a conformational change. This indicates that the transition from the 70S IC to the 70S EC is largely regulated by the release of P*_i_* from IF2 and/or the subsequently rapid release of the GDP-form of IF2 from the 70S IC.

As the 70S IC matures into a 70S EC, dissociation of IF2 disrupts the IF2-ribosome interactions that stabilize the semi-rotated inter-subunit orientation of the 70S IC. Disruption of these IF2-ribosome interactions therefore triggers the reverse rotation of the 30S subunit by ~3° (Figure 3a), which allows the 70S ribosome within the 70S EC to occupy the non-rotated intersubunit orientation. Dissociation of IF2 also disrupts the contact between dIV of IF2 and fMet-tRNA^fMet^, an event that, simultaneously with the reverse rotation of the 30S subunit (at least at our time resolution), enables the central domain and 3’ CCA-fMet tail of fMet-tRNA^fMet^ to move by ~28 Å and ~22 Å, respectively, from the 70S P/I configuration of fMet-tRNA^fMet^ that is observed in the 70S IC to the P/P configuration of fMet-tRNA^fMet^ that is observed in the 70S EC (Figure 3b, 3c, and 3d). This rearrangment of fMet-tRNA^fMet^ is accompanied by an ‘untangling’ of the 3’ CCA-fMet tail that allows the fMet moiety to acquire its peptidyl transfer-competent position within the P site of the PTC (Figure 3c and 3d). Given the simultaneous nature of these conformational changes, at least at our time resolution, we propose that the transition of the 70S ribosome into its to non-rotated inter-subunit orientation is coupled to the rerrangement of fMet-tRNA^fMet^ into its P/P configuration in the 70S EC along with the untangling of the 3’ CCA-fMet tail and positioning of the fMet moiety into the PTC.

**Figure 3:**
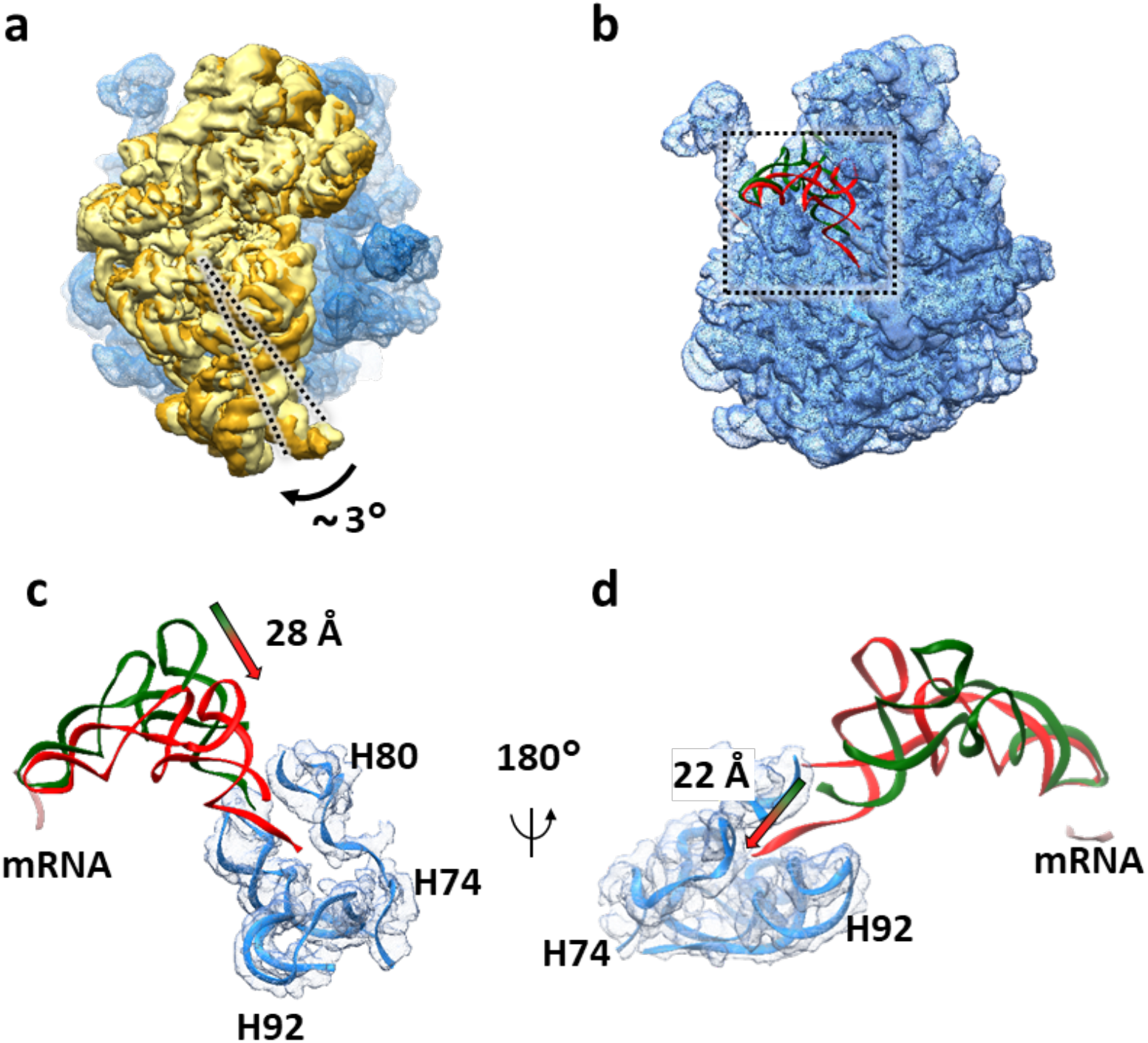
Ribosome and fMet-tRNA^fMet^ dynamics during maturation of the 70S IC into a 70S EC. **(a)** Superposition of the 50S subunits from the 70S IC and 70S EC and comparative analysis of the conformations of the 30S subunit from the 70S IC (pale yellow) and the 30S subunit from the 70S EC (golden yellow). The analysis reveals that the ribosomal subunits from the 70S IC that is initially formed upon 50S subunit joining to the 30S IC transiently acquire a semi-rotated inter-subunit orientation and subsequently undergo an ~3°clockwise rotation, when viewed from the solution side of the 30S subunit, into the non-rotated inter-subunit conformation upon maturation of the 70S IC into a 70S EC. **(b)** Superposition of the 50S subunits from the 70S IC and 70S EC and comparative analysis of the conformations of fMet-tRNA^fMet^ in the 70S P/I configuration from the 70S IC and fMet-tRNA^fMet^ (green) in the P/P configuration from the 70S EC (red). The start codon of the mRNA is shown in pale pink. The analysis reveals the conformational rearrangements of fMet-tRNA^fMet^ that take place as the 70S IC matures into a 70S EC. **(c)** A magnified view of the superposition shown in panel (b) reveals that the central domain of fMet-tRNA^fMet^ moves by ~28 Å towards the P site. **(d)** A 180° rotation of the superposition shown in panel (c) highlights the untangling of the 3’ CCA fMet tail of the fMet-tRNA^fMet^ and its ~22 Å movement into the PTC.

Based on our collective observations, we propose a structure-based model for the late steps of bacterial translation initiation (Figure 4). Within a time range shorter than ~20 ms after mixing 50S subunits and 30S ICs, initial association of 50S subunits to the majority of 30S ICs results in formation of ‘Pre-70S ICs’, that are too transient for us to observe at our current time resolution and that very rapidly either dissociate into free 50S subunits and 30S ICs^13,15,16^ or are converted to 70S ICs. Conversion of the majority of Pre-70S ICs into 70S ICs occurs within ~20-80 ms after mixing 50S subunit and 30S ICs and begins with the rapid hydrolysis of GTP on IF2. Subsequent dissociation of IF1 enables repositioning of dIV of IF2 towards the inter-subunit face of the 30S subunit and formation of the three sets of IF2-ribosome contacts as well as the full constellation of inter-subunit bridges that allow the ribosome to stably occupy the semi-rotated inter-subunit orientation and fMet-tRNA^fMet^ to adopt its 70S P/I configuration. The majority of the resulting 70S ICs mature into 70S ECs within the next several hundreds of ms, a process that begins with release of Pi from IF2 and dissociation of the GDP-form of IF2 from the 70S IC. Dissociation of IF2 enables rotation of the ribosomal subunits into their non-rotated inter-subunit orientation, rearrangement of fMet-tRNA^fMet^ into its P/P configuration, untangling of the 3’ CCA-fMet tail of fMet-tRNA^fMet^, and relocation of the fMet moiety of fMet-tRNA^fMet^ into the PTC, thereby completing 70S EC formation. Notably, we did not observe formation of the minor population of the 70S IC reported by Sprink and coworkers (*i.e.*, 70S-IC I)^9^, suggesting that this conformation of the 70S IC might represent an off-pathway intermediate that is formed only when the 70S IC is trapped when using the GDPNP-form of IF2 and/or prepared using a steady-state approach. In contrast, because the conformation of the 70S IC that we observe here was obtained using the native, GTP-bound form of IF2 under pre-steady-state conditions, we can be certain that it represents a *bona fide* intermediate that is formed on the initiation reaction pathway.

**Figure 4:**
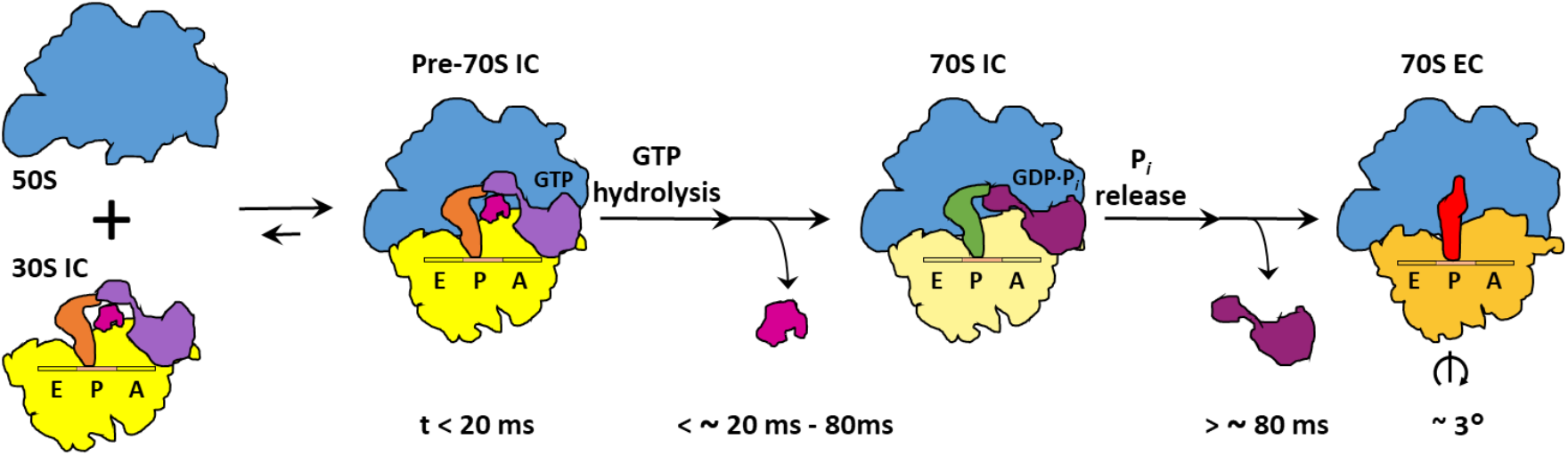
Structure-based kinetic model for late steps in bacterial translation initiation. Cartoon depicting the timing of structural and molecular events during late steps in bacterial translation initiation. Within the first ~20 ms after mixing 50S subunits and 30S ICs, 50S subunits (blue) reversibly join to the majority of 30S ICs (yellow) to form transient Pre-70S ICs. Conversion of the majority of these Pre-70S ICs into 70S ICs takes place within ~20-80 ms after mixing of 50S subunits and 30S ICs and begins with the rapid hydrolysis of GTP on IF2 (light purple). GTP hydrolysis is followed by the dissociation of IF1 (magenta), repositioning of dIV of IF2 (dark purple), and formation of IF2-ribosome interactions and inter-subunit bridges that stabilize the ribosome in its semi-rotated inter-subunit orientation and the fMet-tRNA^fMet^ in its 70S P/I configuration. Within the next several hundred milliseconds, the majority of 70S ICs mature into 70S ECs in a process that begins with release of Pi from IF2 and dissociation of the GDP-form of IF2 from the 70S IC, events that enable rotation of the ribosomal subunits into their non-rotated inter-subunit orientation, rearrangement of fMet-tRNA^fMet^ into its P/P configuration, untangling of the 3’ CCA-fMet tail of fMet-tRNA^fMet^, and relocation of the fMet moiety of fMet-tRNA^fMet^ into the PTC in preparation for formation of the first peptide bond upon delivery of the first aminoacyl-tRNA into the ribosomal aminoacyl-tRNA binding (A) site.

In this report, we have shown how mixing-spraying TR cryo-EM is able to capture physiologically relevant, short-lived, structural intermediates in a biomolecular reaction and have used this approach to elucidate the molecular mechanism of bacterial translation. Although many biophysical and structural biological techniques can operate at higher time resolutions and/or broader timescales than mixing-spraying TR cryo-EM^35^, most of these techniques can only provide local-site or low-spatial-resolution information (*e.g.*, optical spectroscopies) or are otherwise limited to relatively small biomolecular systems of < 50 kDa (*e.g.*, nuclear magnetic resonance spectroscopy). In contrast, mixing-spraying TR cryo-EM is a technique that can provide complete, three-dimensional visualizations of the conformational rearrangements of large biomolecular complexes in real time and at near-atomic spatial resolution. Moreover, because it follows a pre-steady-state reaction that does not need to be inhibited by an analog of a native ligand and/or an inhibitor, one can be certain that the transient conformation(s) observed by mixing-spraying TR cryo-EM are on the reaction pathway. Mixing-spraying TR cryo-EM is a new and powerful structural biology technique that we expect will be employed to follow the formation and maturation of reaction intermediates and elucidate the molecular mechanisms of fundamental biomolecular reactions such as replication, transcription, pre-mRNA processing and splicing, and mRNA and protein degradation.

## METHODS

### Preparation and purification of IC components and assembly of the 30S IC

30S and 50S subunits were purified from the MRE600 *Escherichia coli* strain as previously described, with minor modifications^36^. Tight-coupled 70S ribosomes were isolated by ultracentrifugation of crude ribosomes through a 10–40% sucrose density gradient prepared in Ribosome Storage Buffer (10 mM tris(hydroxymethyl)aminomethane acetate (Tris-OAc) (pH_4 °C_ = 7.5), 60 mM ammonium chloride (NH_4_Cl), 7.5 mM magnesium chloride (MgCl_2_), 0.5 mM ethylenediaminetetraacetic acid (EDTA), 6 mM 2-mercaptoethanol (BME). To maximize the purity of our tight-coupled 70S ribosomes and minimize contaminatination by free 50S subunits, a second round of ultracentrifugation through a 10–40% sucrose density gradient prepared in Ribosome Storage Buffer was added to our standard ribosome purification protocol. Highly pure, tight-coupled, 70S ribosomes were buffer exchanged into Ribosome Dissociation Buffer (10 mM Tris-OAc (pH_4°C_ = 7.5), 60 mM NH_4_Cl, 1 mM MgCl_2_, 0.5 mM EDTA, 6 mM BME) using a centrifugal filtration device (Amicon Ultra, Millipore) with a 100 KDa molecular weight cut off (MWCO) to promote the dissociation of ribosomes into 30S and 50S subunits. 30S and 50S subunits were isolated from the dissociated tight-coupled 70S ribosomes by ultracentrifugation through a 10–40% sucrose density gradient prepared in Ribosome Dissociation Buffer. To ensure high purity, 30S and 50S subunits isolated from the first gradient were subjected to a second round of ultracentrifugation through a 10–40% sucrose density gradient prepared in Ribosome Dissociation Buffer. Highly purified 30S and 50S subunits were concentrated and buffer exchanged into Ribosome Storage Buffer using a centrifugal filtration device with a 100 KDa MWCO. After determining the concentration of the 30S and 50S subunits, small aliquots were prepared, flash frozen in liquid nitrogen, and stored at –80 °C.

The purity of our 30S and 50S subunits was confirmed by negative staining electron microscopy (EM). Briefly, one aliquot of the highly purified 30S subunits was diluted to 50 nM with Tris-Polymix Buffer (50 mM Tris-OAc (pH_RT_ = 7.5), 100 mM potassium chloride (KCl), 5 mM ammonium acetate (NH4OAc), 0.5 mM calcium acetate (CaOAc_2_), 5 mM magnesium acetate (MgOAc_2_), 0.1 mM EDTA, 6 mM BME, 5 mM putrescine dihydrochloride, and 1 mM spermidine, free base). Subsequently, 3 μl of this 50 nM 30S subunit solution was applied to a carbon-coated EM grid for 30 s. Any excess sample solution was wicked away from the EM grid using filter paper, thereby generating a thin layer of sample solution on the EM grid. Following this, 3 μl of a 2 % solution of uranium acetate in water was applied to the EM grid and the EM grid was incubated for 30 s at room temperature. Excess sample solution was wicked away from the EM grid using filter paper, once again generating a thin layer of sample solution on the EM grid. This uranium acetate, negative staining procedure was repeated two more times and, subsequently, the negatively stained 30S subunits were imaged using a 200 kV F20 cryogenic transmission electron microscope (TEM; FEI). Visual inspection of the images that were obtained revealed a highly uniform set of particles exhibiting the characteristically elongated shape of the 30S subunit, thereby demonstrating the purity of the 30S subunits. Analogous procedures were followed to load, negatively stain, and image the highly purified 50S subunits, with visual inspection of the images revealing a highly uniform set of particles exhibiting the characteristic ‘crown view’ of the 50S subunit, thereby demonstrating the purity of the 50S subunits.

IF1 and the γ-isoform of IF2 containing tobacco etch virus (TEV) protease-cleavable, N-terminal, hexa-histidine (6×His) tags were overexpressed in BL21(DE3) cells and purified as described previously^29^. Briefly, 6×His-tagged IFs were purified by nickel nitrilotriacetic acid (Ni^2+^-NTA) affinity chromatography using a batch-binding and elution protocol. After elution of the 6×His-tagged IFs, the 6×His-tags were removed by adding TEV protease to the purified IFs and dialyzing-incubating the mixture overnight (~12 hr) at 4 °C against TEV Cleavage Buffer (20 mM tris(hydroxymethyl)aminomethane hydrochloride (Tris-HCl) (pH4°C = 7.5), 200 mM NaCl, 0.1% Triton X-100, and 2 mM BME). IF1 was further purified on a HiLoad 16/60 Superdex 75 prep grade gel filtration column (GE Biosciences), and IF2 was further purified on a HiTrap SP HP cation-exchange column (GE Biosciences). The purified IFs were concentrated and buffer exchanged into 2× Translation Factor Buffer (20 mM Tris-OAc (pH4°c = 7.5), 100 mM KCl, 20 mM MgOAc2, 10 mM BME) using a centrifugal filtration device (Amicon Ultra, Millipore) with either a 3.5 KDa (IF1) or a 10 KDa (IF2) MWCO. Concentrated IFs were diluted with one volume of 100% glycerol and stored at –20 °C. Prior to using in 30S IC assembly reactions, the IFs were buffer exchanged into Tris-Polymix Buffer using either a Micro Bio-Spin 6 (IF1) or 30 (IF2) gel filtration spin column (Bio-Rad).

tRNA^fMet^ (MP Biomedicals) was aminoacylated and formylated as described previously^29^. The yield of fMet-tRNA^fMet^, which was assessed by hydrophobic interaction chromatography (HIC) on a TSKgel Phenyl-5PW column (Tosoh Bioscience) as described previously^29^, was ~90%. The mRNA used in the 30S IC assembly reaction was chemically synthesized (Thermo Fisher) and is a non-biotinylated variant of the bacteriophage T4 gene product (gp) 32 mRNA that we have used extensively in our single-molecule fluorescence studies of initiation^16,36–38^. The sequence of this mRNA is 5’-CAACCUAAAACUUACACAAAUUAAAAAGGAAAUAGACAUGUUCAAAGUCGAAAAAUCUACUGCU-3’.

The 30S IC was assembled by combining 3.6 μM each of IF1, IF2 and fMet-tRNA^fMet^, 4.8 μM of mRNA, 1 mM GTP, and 2.4 μM of 30S subunits in Tris-Polymix Buffer. The final volume of the 30S IC assembly reaction was 100 μl. IF2, which has been previously shown to protect fMet-tRNA^fMet^ from deacylation^39^ was added to the 30S IC assembly reaction prior to fMet-tRNA^fMet^. To ensure that the 30S IC assembly reaction proceeded in a native, unbiased manner, the 30S subunits were added last. Assembly reactions were incubated at 37 °C for 10 minutes, chilled on ice for 5 minutes, flash frozen in liquid nitrogen and stored at –80 °C.

### Modeling the kinetics of subunit joining and selecting the time points for mixing-spraying TR cryo-EM

To model the kinetics of the subunit joining reaction shown in Extended Figure 2, we used initial 50S subunit and 30S IC concentrations analogous to those used in our mixing-spraying microfluidic chip (*i.e.*, 0.6 μM and 1.2 μM, respectively,), and employed MATLAB to numerically solve a set of differential equations derived from the kinetic scheme and set of rate constants reported by Goyal and coworkers for a subunit joining reaction performed in the presence of IF1 and IF2, but in absence of the IF3^19,31^:

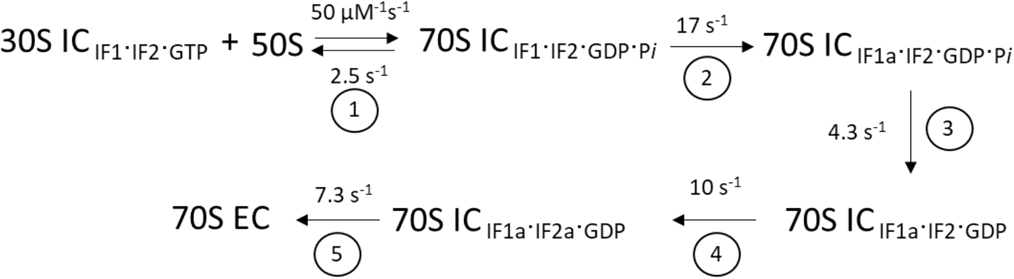

where 30S IC_IF1·IF2·GTP_ is analogous to our 30S IC, 50S is analogous to our 50S subunit, 70S ICif1·if2·gdp·p, is a 70S IC in which GTP has been hydrolyzed to GDP•P*_i_* on IF2, 70S IC_IF1a·IF2·GDP·*P_i_*_ is a 70S IC in which IF1 and/or some other component of the 70S IC has undergone a conformational change, 70S IC_IF1b·IF2·GDP_ is a 70S IC in which P*_i_* has been released from IF2, 70S IC_IF1a·IF2a·GDP_ is a 70S IC in which IF2 has undergone a conformational change, and 70S EC is analogous to our 70S EC. Because of the likelihood that we would not be able to structurally distinguish 70S IC_IF1·IF2·GDP·P*_i_*_, 70S IC_IF1a·IF2·GDP·P_*i*__, 70S IC_IF1a·IF2·GDP_, and 70S IC_IF1a·IF2a·GDP_ at our expected spatial and/or temporal resolutions, we summed the concentrations of these four complexes to generate the grey curve representing the 70S IC in Extended Figure 2. The modeled kinetics predict that the population of 70S ICs should peak within 50–250 ms after mixing of the 50S subunit and 30S IC and that these 70S ICs should mature to a significant population of 70S ECs within the next several hundreds of ms. Thus, to ensure that we would capture formation of the 70S IC and its maturation to the 70S EC, we selected microfluidic chips designed to provide reaction times of ~20 ms, ~80 ms, ~200 ms, and ~600 ms.

### Preparation of EM grids and mixing-spraying TR cryo-EM

Quantifoil R1.2/1.3 grids with a 300 mesh size were subjected to glow discharge in H2 and O2 for 25 s using a Solarus 950 plasma cleaning system (Gatan) set to a power of 25 W. For each of the four time points, 1.2 μM of 50S subunit in T ris-Polymix Buffer and 2.4 μM of 30S IC in Tris-Polymix Buffer were injected into the corresponding microfluidic chip at a rate of 3 μl s^−1^ such that they could be mixed and sprayed onto a glow-discharged grid as previously described^25^. The final concentration of the 50S subunit and the 30S IC after rapid mixing in our microfluidic chip was 0.6 μM and 1.2 μM, respectively. As the mixture was sprayed onto the grid, the grid was plunge-frozen in liquid ethane and stored in liquid nitrogen until it was ready to be imaged.

### Cryo-EM data collection

Plunge-frozen grids were imaged with a 300 kV Tecnai Polara F30 TEM (FEI). The images were recorded at a defocus range of 1-3 μm on a K2 direct detector camera (Gatan) operating in counting mode with an effective magnification of 29,000× at 1.66 Å pixel^−1^. A total of 40 frames were collected with an electron dose of 8 e^−^ pixel^−1^ s^−1^ for each image.

### Cryo-EM data processing

A flow-chart of the data processing procedure detailed here is given in Extended Figure 4. The beam-induced motion of the sample captured by the images was corrected using the MotionCor2 software program^40^. The contrast transfer function (CTF) of each micrograph was estimated using the CTFFIND4 software program^41^. Imaged particles were picked using the Autopicker algorithm included in the RELION 2.0 software program^42^. For each time point, 2D classification of the recorded images was used to separate 30S subunit-like, 50S subunit-like, and 70S ribosome-like particles from ice-like and/or debris-like particles picked by the Autopicker algorithm and to classify the 298K particles that were picked for further analysis into 30S subunit-like, 50S subunit-like, and 70S ribosome-like particle classes. These particle classes were then combined into a single dataset of 30S subunit-like, 50S subunit-like, and 70S ribosome-like particles and subjected to a round of 3D classification using 4× binning of the images to more finely separate 30S subunit-like particles from 50S subunit-like and 70S ribosome-like particles. After the first round of 3D classification, two particle sets were created in which the first particle set encompassed 150K 30S subunit-like particles and the second particle set encompassed 98K combined 50S subunit-like and 70S ribosome-like particles. These two particle sets were extracted using 2× binning of the images.

The first particle set, containing 150K particles, was subjected to a second round of 3D classification from which we then obtained two major subclasses in which the first subclass encompassed 86K 30S ICs and 64K 30S subunits carrying only a P-site fMet-tRNA^fMet^ (or, more likely, a mixture of fMet-tRNA^fMet^ and deacylated tRNA^fMet^) in the 70S P/I configuration. The subclass containing the 30S IC was further refined without binning the images. The resolution of the refined 30S IC was estimated to be 4.2 Å using a resolution-estimating protocol that avoids overfitting and uses the FSC with the 0.143 criterion^32^.

The second particle set, containing 98K combined 50S subunit-like and 70S ribosome-like particles, was also subjected to a second round of 3D classification from which we then obtained two major subclasses in which the first subclass encompassed 21K 50S subunits and the second subclass encompassed 77K 70S ribosome-like particles. This second round of 3D classification was repeated three times to estimate the errors associated with the 3D classification-based division of the 50S subunit and 70S ribosome-like particle populations (Table 1). At this point in the analysis, each 50S subunit and 70S ribosome-like particle was traced back to the time point from which it originated to determine the 50S subunit and 70S ribosome-like particle populations at each time point. The 70S ribosome-like subclass was then subjected to a third round of 3D classification from which we then obtained two major sub-subclasses in which the first sub-subclass encompassed 27K 70S ICs and the second sub-subclass encompassed 50K 70S ECs.

**Table 1.** Populations of the 50S subunit, 70S IC, and 70S EC obtained after 3D classification. Standard deviations were obtained by repeating the 3D classification procedure three times for each time point.

|  | 50S (%) | 70S IC (%) | 70S EC (%) |
| --- | --- | --- | --- |
| **20 ms** | 64.6 ± 5.8 | 18.3 ± 0.5 | 17.1 ± 0.5 |
| **80 ms** | 20.1 ± 1.8 | 67.2 ± 0.6 | 12.7 ± 0.6 |
| **200 ms** | 18.0 ± 3.7 | 49.8 ± 7.6 | 32.2 ± 7.6 |
| **600 ms** | 14.7 ± 1.4 | 25.1 ± 6.7 | 60.3 ± 6.7 |

Again, this third round of 3D classification was repeated three times in order to estimate the errors associated with the 3D classification-based division of the 70S IC and 70S EC populations (Table 1). The subclass containing the 50S subunit and the sub-subclasses containing the 70S IC and 70S EC were then further refined without binning the images. The resolutions of the refined 70S IC and 70S EC were estimated to be 4.0 Å and 3.9 Å, respectively, using a resolution-estimating protocol that avoids overfitting and uses the FSC with the 0.143 criterion^32^.

### Assessing the integrity of the 30S IC during mixing-spraying TR cryo-EM

To assess whether the 30S IC was stable enough to maintain its integrity during mixing-spraying TR cryo-EM, 2.4 μM of 30S IC in Tris-Polymix Buffer and an equal volume of Tris-Polymix Buffer lacking 50S subunits were injected into the microfluidic chip designed to give the longest reaction time (~600 ms), mixed, and sprayed onto an EM grid as the grid was plunge-frozen in liquid ethane and subsequently stored in liquid nitrogen, all as described above. When ready, the plunge-frozen grid was imaged with the 300 kV Tecnai Polara F30 TEM (FEI), as described above. Subsequently, 2D classification was used to select 30S subunit-like particles. The selected particles were then subjected to 3D classification, which showed that ~75% of the 30S subunit-like particles were 30S ICs and ~25 % of the 30S subunit-like particles were 30S subunits carrying a P-site fMet-tRNA^fMet^ (or, more likely, a mixture of fMet-tRNA^fMet^ and deacylated tRNA^fMet^) in its ‘30S P/I’ configuration (Extended Figure 1). The results of this experiment demonstrate that the majority of the 30S IC remains intact during injection and mixing in the microfluidic chip and spraying onto the EM grid.

### Modeling of the 30S IC, 70S IC, and 70S EC structures

We obtained near-atomic resolution models of the 30S IC and 70S IC by employing the Molecular Dynamics Flexible Fitting (MDFF) method^33^ (Extended Figure 8) and using a near-atomic-resolution cryo-EM model of a 70S IC (PDB ID: 3JCJ) as the initial starting model.

Similarly, we obtained an initial, near-atomic resolution model of the 70S EC using rigid-body fitting within the UCSF Chimera software program^43^ and atomic-resolution models of 70S ribosomes in the non-rotated inter-subunit orientation and lacking any tRNA or mRNA ligands (PDB IDs: 2AVY and 2AW4). This initial, near-atomic-resolution model of the 70S EC was further refined by subjecting it to the ‘jiggle fit’ algorithm within the COOT software program^44^ to obtain the final atomic coordinates.

## DATA AVAILABILITY

The data that support the findings of this study are available from the corresponding author upon request.

## ACKNOWLEDGEMENTS

This work was supported by funds to J.F. from the National Institutes of Health (R01 GM 55440 and GM 29169) and to R.L.G. from the National Institutes of Health (R01 GM 084288). K.C. was supported by an American Cancer Society Postdoctoral Fellowship (125201).

## AUTHOR CONTRIBUTIONS

S.K., Z.F., K.C., B.C., M.S., R.L.G., and J.F. designed the research; K.C. prepared all of the biological reagents; S.K. and Z.F. performed the time-resolved cryo-EM experiments; S.K., Z.F., and W.L. analyzed the data; S.K., K.C., R.L.G., and J.F. wrote the manuscript; all eight authors approved the final manuscript.

## REFERENCES

1. Antoun, A., Pavlov, M.Y., Andersson, K., Tenson, T. & Ehrenberg, M. The roles of initiation factor 2 and guanosine triphosphate in initiation of protein synthesis. Embo Journal 22, 5593–5601 (2003).

2. Hussain, T., Llacer, J.L., Wimberly, B.T., Kieft, J.S. & Ramakrishnan, V. Large-Scale Movements of IF3 and tRNA during Bacterial Translation Initiation. Cell 167, 133-+ (2016).

3. Julian, P. et al. The Cryo-EM Structure of a Complete 30S Translation Initiation Complex from Escherichia coli. Plos Biology 9 (2011).

4. Lopez-Alonso, J.P. et al. Structure of a 30S pre-initiation complex stalled by GE81112 reveals structural parallels in bacterial and eukaryotic protein synthesis initiation pathways. Nucleic Acids Research 45, 2179–2187 (2017).

5. Simonetti, A. et al. Involvement of protein IF2 N domain in ribosomal subunit joining revealed from architecture and function of the full-length initiation factor. Proceedings of the National Academy of Sciences of the United States of America 110, 15656–15661 (2013).

6. Simonetti, A. et al. Structure of the 30S translation initiation complex. Nature 455, 416–U73 (2008).

7. Allen, G.S., Zavialov, A., Gursky, R., Ehrenberg, M. & Frank, J. The cryo-EM structure of a translation initiation complex from Escherichia coli. Cell 121, 703–712 (2005).

8. Myasnikov, A.G. et al. Conformational transition of initiation factor 2 from the GTP- to GDP-bound state visualized on the ribosome. Nature Structural & Molecular Biology 12, 1145–1149 (2005).

9. Sprink, T. et al. Structures of ribosome-bound initiation factor 2 reveal the mechanism of subunit association. Science Advances 2 (2016).

10. Frank, J. Time-resolved cryo-electron microscopy: Recent progress. Journal of Structural Biology 200, 303–306 (2017).

11. Antoun, A., Pavlov, M.Y., Lovmar, M. & Ehrenberg, M. How initiation factors maximize the accuracy of tRNA selection in initiation of bacterial protein synthesis. Molecular Cell 23, 183–193 (2006).

12. Caban, K. & Gonzalez, R.L. The emerging role of rectified thermal fluctuations in initiator aa-tRNA- and start codon selection during translation initiation. Biochimie 114, 30–38 (2015).

13. Milon, P., Konevega, A.L., Gualerzi, C.O. & Rodnina, M.V. Kinetic checkpoint at a late step in translation initiation. Molecular Cell 30, 712–720 (2008).

14. Milon, P. & Rodnina, M.V. Kinetic control of translation initiation in bacteria. Critical Reviews in Biochemistry and Molecular Biology 47, 334–348 (2012).

15. Grigoriadou, C., Marzi, S., Kirillov, S., Gualerzi, C.O. & Cooperman, B.S. A quantitative kinetic scheme for 70 S translation initiation complex formation. Journal of Molecular Biology 373, 562–572 (2007).

16. MacDougall, D.D. & Gonzalez, R.L. Translation Initiation Factor 3 Regulates Switching between Different Modes of Ribosomal Subunit Joining. Journal of Molecular Biology 427, 1801–1818 (2015).

17. Gualerzi, C.O. & Pon, C.L. Initiation of mRNA translation in bacteria: structural and dynamic aspects. Cellular and Molecular Life Sciences 72, 4341–4367 (2015).

18. Wilson, D.N. Ribosome-targeting antibiotics and mechanisms of bacterial resistance. Nature Reviews Microbiology 12, 35–48 (2014).

19. Goyal, A., Belardinelli, R., Maracci, C., Milon, P. & Rodnina, M.V. Directional transition from initiation to elongation in bacterial translation. Nucleic Acids Research 43 (2015).

20. Huang, C.H., Mandava, C.S. & Sanyal, S. The Ribosomal Stalk Plays a Key Role in IF2-Mediated Association of the Ribosomal Subunits. Journal of Molecular Biology 399, 145–153 (2010).

21. Tomsic, J. et al. Late events of translation initiation in bacteria: a kinetic analysis. Embo Journal 19, 2127–2136 (2000).

22. Ling, C. & Ermolenko, D.N. Initiation factor 2 stabilizes the ribosome in a semirotated conformation. Proceedings of the National Academy of Sciences of the United States of America 112, 15874–15879 (2015).

23. Marshall, R.A., Aitken, C.E. & Puglisi, J.D. GTP Hydrolysis by IF2 Guides Progression of the Ribosome into Elongation. Molecular Cell 35, 37–47 (2009).

24. LaTeana, A., Pon, C.L. & Gualerzi, C.O. Late events in translation initiation. Adjustment of fMet-tRNA in the ribosomal P-site. Journal of Molecular Biology 256, 667–675 (1996).

25. Chen, B. et al. Structural Dynamics of Ribosome Subunit Association Studied by Mixing-Spraying Time-Resolved Cryogenic Electron Microscopy. Structure 23, 1097–1105 (2015).

26. Fu, Z. et al. Key Intermediates in Ribosome Recycling Visualized by Time-Resolved Cryoelectron Microscopy. Structure 24, 2092–2101 (2016).

27. Lu, Z. et al. Monolithic microfluidic mixing-spraying devices for time-resolved cryo-electron microscopy. Journal of Structural Biology 168, 388–395 (2009).

28. Shaikh, T.R. et al. Initial bridges between two ribosomal subunits are formed within 9.4 milliseconds, as studied by time-resolved cryo-EM. Proceedings of the National Academy of Sciences (2014).

29. Fei, J.Y. et al. A Highly Purified, Fluorescently Labeled in Vitro Translation System for Single-Molecule Studies of Protein Synthesis. Methods in Enzymology, Vol 472: Single Molecule Tools, Pt A: Fluorescence Based Approaches 472, 221–259 (2010).

30. Grigoriadou, C., Marzi, S., Pan, D., Gualerzi, C.O. & Cooperman, B.S. The translational fidelity function of IF3 during transition from the 30 S initiation complex to the 70 S initiation complex. Journal of Molecular Biology 373, 551–561 (2007).

31. Goyal, A. Monitoring the late events of translation initiation in real-time, Doctoral Dissertation, (Georg-August University Göttingen, Göttingen, 2015).

32. Chen, S.X. et al. High-resolution noise substitution to measure overfitting and validate resolution in 3D structure determination by single particle electron cryomicroscopy. Ultramicroscopy 135, 24–35 (2013).

33. Trabuco, L.G., Villa, E., Mitra, K., Frank, J. & Schulten, K. Flexible fitting of atomic structures into electron microscopy maps using molecular dynamics. Structure 16, 673–683 (2008).

34. Wang, J., Liu, Z., Frank, J. & Moore, P.B. Identification of ions in experimental electrostatic potential maps. IUCrJ 5, 375–381 (2018).

35. Henzler-Wildman, K. & Kern, D. Dynamic personalities of proteins. Nature 450, 964–972 (2007).

36. Caban, K., Pavlov, M., Ehrenberg, M. & Gonzalez, R.L. A conformational switch in initiation factor 2 controls the fidelity of translation initiation in bacteria. Nature Communications 8 (2017).

37. Wang, J., Caban, K. & Gonzalez, R.L., Jr. Ribosomal initiation complex-driven changes in the stability and dynamics of initiation factor 2 regulate the fidelity of translation initiation. J Mol Biol 427, 1819–34 (2015).

38. Elvekrog, M.M. & Gonzalez, R.L., Jr. Conformational selection of translation initiation factor 3 signals proper substrate selection. Nat Struct Mol Biol 20, 628–33 (2013).

39. Guenneugues, M. et al. Mapping the fMet-tRNA(f)(Met) binding site of initiation factor IF2. EMBO J 19, 5233–40 (2000).

40. Zheng, S.Q. et al. MotionCor2: anisotropic correction of beam-induced motion for improved cryo-electron microscopy. Nature Methods 14, 331–332 (2017).

41. Rohou, A. & Grigorieff, N. CTFFIND4: Fast and accurate defocus estimation from electron micrographs. Journal of Structural Biology 192, 216–221 (2015).

42. Scheres, S.H.W. RELION: Implementation of a Bayesian approach to cryo-EM structure determination. Journal of Structural Biology 180, 519–530 (2012).

43. Pettersen, E.F. et al. UCSF chimera - A visualization system for exploratory research and analysis. Journal of Computational Chemistry 25, 1605–1612 (2004).

44. Emsley, P. & Cowtan, K. Coot: model-building tools for molecular graphics. Acta Crystallographica Section D-Biological Crystallography 60, 2126–2132 (2004).

